# SARS-CoV-2 spike proteins uptake mediated by lipid raft ganglioside GM1 in human cerebrovascular cells

**DOI:** 10.1101/2022.03.20.485050

**Authors:** Conor McQuaid, Alexander Solorzano, Ian Dickerson, Rashid Deane

**Affiliations:** Del Monte Institute of Neuroscience, Department of Neuroscience, University of Rochester, URMC, 601 Elmwood Avenue, Rochester, NY 14642

**Keywords:** Brain endothelial cells, pericytes, vascular smooth muscle cells, COVID-19 Neurological symptoms, SARS-CoV-2 variants of interest, caveolin

## Abstract

While there is clinical evidence of neurological manifestation in coronavirus disease-19, it’s unclear whether this is due to differential severe acute respiratory syndrome coronavirus 2 (SARS-CoV-2) uptake from blood by cells of the cerebrovasculature. SARS-CoV-2 and its spike protein (SP) interact with the endothelium but the roles of extracellular peptidase domain on angiotensin converting enzyme 2 receptors (ACE2) and ACE2 independent pathways (such as glycans) are not fully elucidated. In addition, for SARS-CoV-2 to enter the brain parenchyma from blood it has to cross several cell types, including the endothelium, pericytes and vascular smooth muscle. Since SARS-CoV-2 interacts with host cells via it SP at the entry point of it life cycle, we used fluorescently labelled SP (SP-555) (wild type and mutants) to model viral behaviour, *in vitro*, for these cell types (endothelial, pericytes and vascular smooth muscle) to explore pathways of viral entry into brain from blood. There was differential SP uptake by these cell types. The endothelial cells had the least uptake, which may limit SP uptake into brain from blood. Uptake was mediated by ACE2, but it was dependent on SP interaction with ganglioside GM1 in the lipid raft. Mutation sites, N501Yand E484K and D614G, as seen in variants of interest, were differentially taken up by these cell types. There was greater uptake but neutralization with anti-ACE2 and anti-GM1antibodies was less effective. Our data suggested that GM1/lipid raft is an important entry point of SARS-CoV-2 into these cells since inhibition of SP uptake with both anti-ACE2 and anti-GM1 together was similar to that with only anti-GM1, and both ACE2 and GM1 are within the lipid raft region of plasma membrane. Thus, GM1 is a potential SARS-CoV-2 and therapeutic target at the cerebrovasculature.

## Introduction

While severe acute respiratory syndrome coronavirus 2 (SARS-CoV-2) primarily elicits respiratory infectious coronavirus disease-19 (COVID-19) (Laprise et al., 2019), many non-respiratory organs are also affected, including the brain (Brady et al, 2021; Huang et al., 2020; L. Mao et al., 2020; McQuaid et al., 2021; Moriguchi et al., 2020; Saleki et al., 2020), heart (Dhakal et al., 2020; Perez-Bermejo et al., 2021; Puntmann et al., 2020), kidneys (Fanelli et al., 2020; Martinez-Rojas et al., 2020), and liver (R. Mao et al., 2020; Marjot et al., 2021; Wang et al., 2020; Zhong et al., 2020). This may suggest that there is viral distribution from the blood. There is continuous evolution of SARS-CoV-2 variants, despite the rapid development of vaccines and therapeutics, that affect humans of all ages with short-and long-term consequences, including neurological manifestations. Since SARS-CoV-2 would need to cross several cell types to enter brain parenchyma, we used three cell types of the human cerebrovasculature (endothelium, pericytes and smooth muscle cells) to determine if there are differential mechanisms of SARS-CoV-2 uptake so as to provide a mechanistic basis for a more targeted therapy.

The spike protein (SP) of SARS-CoV-2 is a structural protein, which is assemble as a trimer of the heterodimer (S1-S2), that protrudes from the viral surface to give it the crown-like appearance (Gordon et al., 2020; Li et al., 2020). The S1 unit contains a receptor binding domain (RBD), which promotes attachment to host cells via facilitators, such as to the extracellular peptidase domain on angiotensin converting enzyme 2 receptors (ACE2) (Doobay et al., 2007; Hamming et al., 2004; Hoffmann et al., 2020; Perrotta et al., 2020; Xu et al., 2020). TMPRESS 2 (transmembrane protease, serine 2) on the host cells cleaves the SP to promote viral entry to cells (Matsuyama et al., 2020).There are other receptors/facilitators on the cell surface that mediate the entry of SARS-CoV-2, including oligosaccharide receptors via sialic acid (Chen et al., 2005; Erickson et al., 2021; Rhea et al., 2021; Tortorici et al., 2019). Thus, there are multiple interaction sites between SARS-CoV-2 and host cells, which may contribute to cell type specific effects and the diverse symptoms of COVID-19 (McQuaid et al., 2021).

SARS-CoV-2 has been detected in brains of severely infected deceased people, however it is unclear as to how it gets there and if this leads to significant viral neuro-invasion (McQuaid et al., 2021). Recombinant spike proteins have been used to study viral behavior by using *in vitro* models of brain endothelial cells, and in vivo studies (Brady et al., 2021; Buzhdygan et al., 2020; Rhea et al., 2021). While it was reported that SP interacts with the brain endothelial cells, this was independent of ACE2, in mice (Rhea et al., 2021). The entry of SARS-CoV-2 into brain requires that it crosses several cell types, including the endothelium at the interface of blood and brain, pericytes and smooth muscles. However, SARS-CoV-2 or SP uptake by these cell types was not characterized.

Herein, we used fluorescently labeled SP of wild type (WT) and mutants (from variants of concern) to establish the mechanism of SP uptake by cerebrovascular cells (endothelial cells, pericytes and smooth muscle cells). We show that there was differential SP uptake by these three cell types, with the endothelial cell, at the interface between blood and brain, showing the lowest capacity for SP uptake. While SP uptake was mediated by ACE2, this was dependent on its interaction with gangliosides, especially monosialotetrahexasylganglioside, GM1, which is present in the lipid raft/caveolin regions of the plasma membrane as well as ACE2. Mutant SPs, N501Y and E484K and D614G, showed a higher uptake compared to control wild type SP, except for D614G in pericytes. Sticking differences were greater binding to sialic acid via wheat germ agglutinin (WGA) and neutralization of the mutant uptake was less effective than that of the wild type SP, using anti-ACE2 and anti-GM1 antibodies. While there were multiple potential SP binding sites on these cells, sialic acid on GM1 plays a major role in SP uptake. Thus, the lipid raft, which also has a high concentration of ACE2, may serve as an important entry point to host cells. By deduction, our data suggested that GM1 is a potential SARS-CoV-2 target, which may lead to addition therapeutics to alleviate some of the diverse neurological and COVID-19 symptoms.

## Materials

SARS-CoV-2 Spike proteins (recombinant SARS-CoV-2 Spike Protein (SP-RBD, Arg319-Phe 541; cat# RP-87678, HEK293 cell expressed and binds ACE2) was obtained from Life Technologies Corporation, Carlsbad CA, USA. Mutants SPs and its control wild type protein were obtained from RayBiotech Inc, Peachtree Corners, GA, USA. Recombinant mutants N501Y (cat# 230-30184, expressed region Arg319-Phe541), D614D (Cat# 230-3030186, expressed region Arg319-Gln690), E484K (cat# 230-30188, expressed region Arg319-Phe541) and their SP (Cat# 230-30162, expressed region Arg319-Phe541) were also HEK 293 expressed. All SPs were labeled separately with Alexa Fluor 555, using a kit (Microscale protein labeling kit; ThermoFisher Scientific; Waltham, MA, USA) and by following the manufacturer instructions. Anti-ACE2 antibody (R&D Systems, Cat# AF933) was labeled with Alexa Fluor 488 by following the manufacturer instructions (Microscale protein labeling kit; ThermoFisher Scientific). In addition, the labeled SPs or anti-ACE2 antibody were purified using 3 kDa molecular weight cut-off ultrafiltration filter (Amicon Ultra Centrifugal Filter, Millipore). There was no detectable dye in the filtrate.

Antibodies raised against the extracellular domain of potential SP binding receptors were used. These include, anti ACE2 antibody (R&D Systems, Cat# AF933) used at a low (10 μg/m)l and high (60 μg/ml) concentration; anti TMPRSSE2 antibody (Invitrogen, Cat# PA5-14264) used at 13 μg/ml; anti CD147 antibody (Invitrogen, Cat# 34-5600) used at 2.5 μg/ml; anti NP-1antibody (Invitrogen, Cat# PA5-47027) used at 2 μg/ml and anti GM1 antibody (Abcam, Cat# Ab23943) used at 5 μg/ml. The concentration used was obtained from the manufacturer guidance. Wheat Germ agglutinin (WGA; Cat# L9640) and heparin (cat# H3393) were obtained from Sigma, and used at 10 μg/mL and 100 μg/mL, respectively. Transferrin from human serum conjugated to Aexa Fluor 488(Cat# T13342), BODIPY FL C5-Lactosylceramide complex to BSA (Cat# B34402) and BODIPY FL ganglioside(Cat# B13950) were obtained from ThermoFisher Scientific) and used at 10 μg/ml. Nystatin (Cat# J62486.09, and chlorpromazine (Cat# J63659) were obtained from ThermoFisher Scientific, and used at 25 μg/ml and 10 μg/ml, respectively.

## Methods

### Cell culture

Human Cerebral Microvascular Endothelial Cells (hCMEC/D3) were purchased from Millipore (#SCC066) and expanded in EBM-2 Endothelial Cell Growth Basal Medium (Lonza #00190860) with EGM-2 MV* Microvascular Endothelial Singlequot kit (Lonza #CC-4147) supplemented media. hCMEC/D3 cells were expanded in T25 flask (ThermoFisher Scientific #156367) on a collagen IV (50 μg/ml Sigma-Aldrich #122-20) based growth matrix. *hCMEC/D3 cells were cultured in modified EBM-2 MV medium (Lonza) containing (v/v) 0.025% VEGF, IGF and EGF, 0.1% bFGF, 0.1% rhFGF, 0.1% gentamycin, 0.1% ascorbic acid and 0.04% hydrocortisone, and 1% 100 U/ml penicillin and 100μg/ml streptomycin. When hCMEC/D3 cells were grown on glass slides (ibidi u-chamber 12 well glass slides #81201) for experiments, collagen and fibronectin (50 μg/ml Millipore FC014) were used as matrix. Human Brain Vascular Smooth Muscle Cells (HBVSMCs) were purchased from Sciencell Research Laboratories, Inc., Carlsbad, CA, USA (Cat#1100) and expanded in Smooth Muscle Cell Medium (Sciencell #1101) by following manufacturer’s guidelines. HBVSMCs cell were expanded in T25 flasks with a Poly-L-Lysine (PLL (0.01% Sigma-Aldrich #P4707)) based growth matrix. When HBVSMCs were grown on glass slides (ibidi u-chamber 12 well glass slides) for experiments, fibronectin (50 μg/ml Millipore FC014) was used as matrix. Human Brain Vascular Pericytes (HBVPs) were purchased from Sciencell (#1200) and expanded in Smooth Muscle Cell Medium (Sciencell #1201) by following manufacturer’s guidelines. HBVSMCs cell were expanded in T25 flasks with a PLL (0.01% Sigma-Aldrich #P4707) based growth matrix. When HBVSMCs were grown on glass slides (ibidi u-chamber 12 well glass slides) for experiments, fibronectin (50 μg/ml Millipore FC014) was used as matrix. All cells were split at 90% confluency into fresh matrix coated flasks or slides for growth and experiments. Medium was changed within 24 hrs of initial split and every 2-3 days thereafter. Cells were kept in an incubator (37°C, 5% CO_2_ humidified) during growth and experiments. Studies at 4°C were performed in a fridge.

### Uptake experiments

Cells were grown to confluent on glass-wells slides (ibidi u-chamber 12-well glass slides), media removed, cells washed 3x with Hank’s Balance Salt Solution with Calcium and Magnesium (HBSS +Ca/Mg, glucose and bicarbonate (Gibco, Waltham, MA, USA 14025-092)) before exposing them to SP diluted in HBSS +Ca/Mg at a given concentration (usually 100 nM) and kept in an incubator (37°C, 5% CO_2_ humidified) for the duration of the experiment. At the end of the experiment, cells were washed with HBSS +Ca/Mg, fixed in 4 % paraformaldehyde (PFA) for 10 mins and mounted with ProLong™ Glass Antifade Mountant with NucBlue™ Stain (ThermoFisher Scientific P36985).

### Inhibition studies

Cells were pre-incubate with the inhibitor (diluted in HBSS) for 15 mins and present with the SP(100 nM) for 4hrs at 37°C, 5% CO_2_ in the humidified incubator. All cells were then washed, fixed, mounted and imaged. Nystatin (Cat# J62486.09, and chlorpromazine (Cat# J63659) were obtained from ThermoFisher Scientific, and used at 25 μg/ml and 10 μg/ml, respectively. Values were calculated as percentage of control wells which were SP uptake without the inhibitor but with the vehicle solution.

### Low temperature studies

Cells were pre-incubate in the fridge (4°C) for 30 mins to adapt the cells to this temperature before running the experiment for 1 hr in the fridge in the presence of SP-555 (100 nM). For comparison, cells were incubated in the incubator for 30 mins followed by 1 hr in the presence of SP-555 (100 nM). All cells were then washed, fixed, mounted and imaged.

### Imaging and analysis

Whole well images were taken on an Olympus VS-120 slide scanner. Exposure and gain were fixed for all experimental groups based on pilot experiments. All fluorescence (SP-555 and DAPI) quantification was performed without enhancement of signals. The person performing the analysis were blinded to the experimental design and tracer used. The VSI files were imported into Qupath for analysis. Five square fields with an area of 1×10^−6^ μm were randomly placed on each imaged. Custom pixel classifiers were created to measure the fluorescence intensity in the red (555 nm) channel. The settings for the pixel classifiers were standardized to be: random trees, moderate pixel resolution, and local mean subtraction set at 1. The cell count was performed with Qupath’s cell counter, the settings varied amongst different cell types. Custom object classifiers for cell detection were created to correct the count of cells in the blue (359 nm) channel. The analysis process was automated using scripts generated by Java Groovy in Qupath. Data were expressed as fluorescence intensity per cell for standardization. Data for figure 1 and 2 were obtained using images from a Axiovert 25 Zeiss Inverted microscope (Obrekochen, Germany). These were not compared with any of the other data.

**Figure 1.**
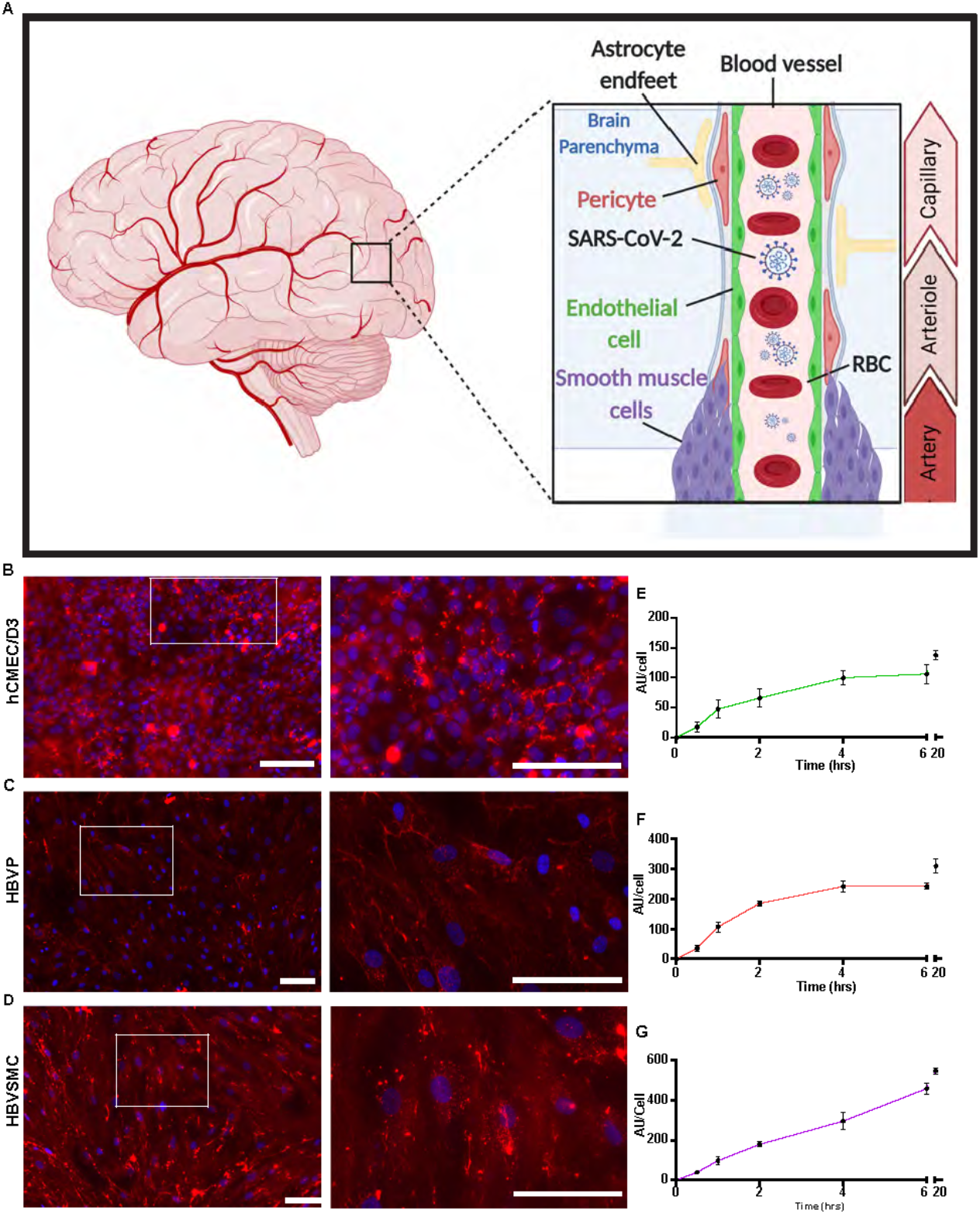
Progressive increase in SP uptake by cerebrovascular cells. **A**. Schematic diagram showing the location of the three human cerebrovascular cell types used in this study (micro vessel/capillaries endothelial cells (hCMEC/D3), pericytes (HBVP) and vascular smooth muscle cells (HBVSMC)). **B-D**. Representative images of SARS-CoV-2 spike protein (SP)-555 (red) uptake, at 4 hrs, counter stained with DAPI. Images on the right is the white boxed areas. SP-555 uptake over time for the hCMEC/D3 (**E**), HBVP (**F**) and HBVSMC (**G**). AU= Fluorescence arbitrary unit. Values are mean ± SEM. N= 3 to 6 wells per group. Scale bar =100 μm.

**Figure 2.**
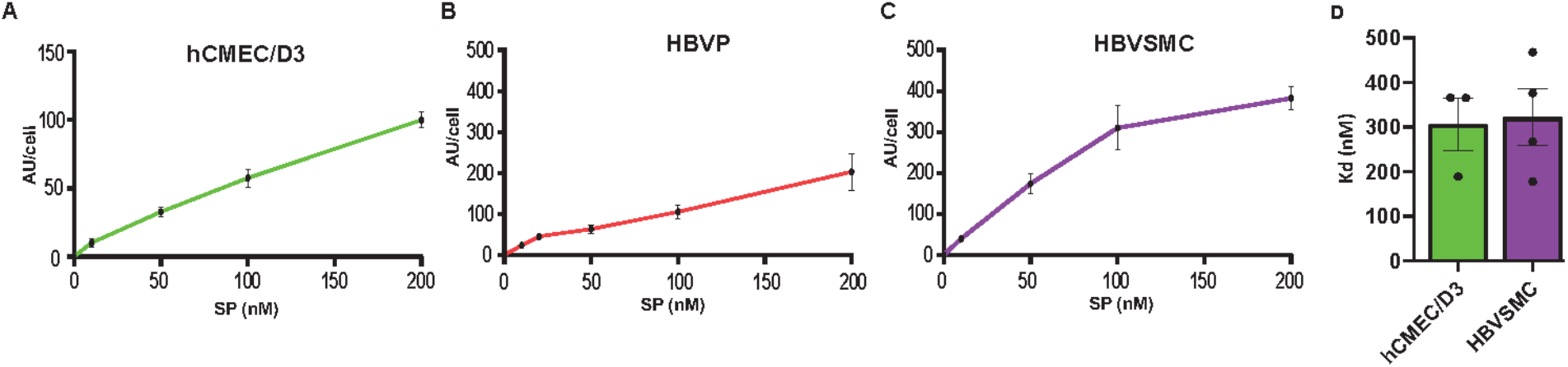
Saturable SP uptake. **A**. SP-555 uptake vs SP-555 concentration for the hCMEC/D3 (**A**), HBVP (**B**) and HBVSMC (**C**). Binding constant (Kd) for the cell types that were best fitted to a saturable uptake. Values are mean **±** SEM. N=3-5 wells/group Au= Fluorescence arbitrary unit. Supplementary Figure 1 and 2.

### Immunocytochemistry (ICC) and imaging

Cells were grown to confluency on glass slides, medium removed, washed with HBSS +Ca/Mg and fixed in 4% PFA for 15mins. Cells were not permeabilized. The same primary antibodies used for inhibition assay were used in the ICC and at the same concentration. The secondary antibodies, which were conjugated to Alexa Fluor 488, were donkey anti-rabbit (Thermofisher Scientific #A32790), anti-goat (Thermofisher Scientific #A32814) and anti-mouse (Thermofisher Scientific #A32766), and used at 1:200 dilution. Cells were mounted using with ProLong™ Glass Antifade Mountant with NucBlue™ Stain (ThermoFisher Scientific P36985) and imaged on Axiovert 25 Zeiss Inverted microscope (Obrekochen, Germany).

### Cell viability

Unlabeled wild type SP (0-200 nM) was used to measure *in vitro* cell viability by the MTT (3-(4,5-dimethylthiazol-2-yl)-2,5-diphenyltetrazolium bromide) assay (Roche, Cell proliferation Kit 1 (MTT) Cat no. 11465007001), a colorimetric method according to which a tetrazolium-based compound is reduced to formazan by living cells. The amount of formazan produced is directly proportional to the number of living cells in the culture. Each cell type was incubated for 24hr at 37C with 5% CO_2_

### ACE2 binding to SP

Recombinant human ACE2 (HEC 293 cells; cat# 230-30165), RayBiotech Life, Inc., Peachtree Corners, GA), dissolved in carbonate/bicarbonate buffer, was immobilized (2 μg/ml) on glass slides for 1 hr at room temperature (RT), blocked with a non-protein buffer (Pierce Blocking buffer), washed, incubated with SP-555 at different concentrations in HBSS for 1 hr at RT, washed, mounted and imaged. SP-555 intensity for each concentration were analysed and expressed as intensity/ μm^2^. Values are mean ± SEM.

### Statistic

All analysis were performed using Graphpad Prims version 9.2.0. Statistically analyzed was by analysis of variance (ANOVA) followed by Tukey post hoc test for three or more groups. Unpaired t-test was used to compared two groups. The differences were considered to be significant at p < 0.05. For statistical representation, *P < 0.05, **P < 0.01, and ***P < 0.001 and ****P< 0.0001 are the levels of statistical significance. All values were expressed as mean ± SEM. N is the number of cell culture wells per group. For each well the average intensity of the five fields was used. Outliers were identified and removed using ROUT method with Q= 10% in GraphPad Prism version 9.2.0.

## Results

### Progressive SARS-CoV-2/SP uptake by cerebrovascular cells

The human brain cerebrovascular cells, endothelial cell (hCMEC/D3), primary pericyte (HBVP) and primary vascular smooth muscle cell (HBVSMC) were used (**Figure 1A**), as a monolayer to characterize the uptake mechanisms of WT spike proteins (SP-555), and mutants that are commonly seen in variants of interest. While SP-555 signal was associated with each cell type, it was mostly seen on the cell surface of hCMEC/D3, but for the other two cell types there were more internalization (**Figure 1B-D**). The time-dependent uptake pattern, determined at 100 nM SP-555 and over 20 hrs, showed that each cell type approached equilibrium after 6 hrs (**Figure 1E-F**) with a half-time (t1/2) of 2 to 3 hrs to equilibrate. The endothelial cells had the lowest capacity to take up SP compared to the other two cell types, possibly due to difference in the cell size.

### Saturable SARS-CoV-2/SP uptake by these cerebrovascular cells

The concentration-dependent SP-555 uptake was determined at 4 hrs, and this showed a pattern of approaching saturation after about 200 nM for the hCMEC/D3 and HBVSMC, while for the HBVP it was linear (**Figure 2A-C**). Higher concentrations were not used as previous studies used <100 nM (Buzhdygan et al, 2020; Rhea et al,2021), and the relevant of higher concentration maybe questionable. The estimated binding affinity was similar for the hCMEC/D3 cells and HBVMC (**Figure 2D**). However, these values were greater than that for ACE2/SP-555 binding, in vitro (**Supplementary Figure 1**), which is similar to that of the manufacturer value. Thus, possible mechanisms are likely similar for these cell types and may be due to multiple uptake mechanisms. We confirm that SP was not toxic to these cell types (**Supplementary Figure 2**). Even though there are limitations in using the MTT assay (Ghasemi et al., 2021), we use the same conditions for each cell type and followed the manufacturer instructions. This assay is dependent on mitochondria metabolism to reduce MTT to formazan, a water soluble violet-blue compound. Thus, the increased levels seen at the higher SP concentration for the hCMEC/D3 may be due to increased metabolic activity due to the higher mitochondria content of cerebral endothelium (Andrade Silva et al., 2021; Mullen et al., 2021; Oldendorf et al., 1977; Oldendorf and Brown, 1975). This should not significantly affect our data since the experiments were performed at or less than 4 hrs, while the MTT assay was performed after 24 hrs incubation with the SP.

### ACE2 mediates SARS-CoV-2/SP uptake

The pattern of SP uptake indicates some receptor binding and these cell types express a number of receptors that are associated with SARS-CoV-2, such as ACE2, a major binding site of SP (**Supplementary Figure 3**). ACE2 is expressed in human brain endothelial cells (Qiao et al., 2020) and in pericytes and vascular smooth muscle cells (He et al., 2020). We confirmed that ACE2 interacts with SP-555, in vitro (**Supplementary Figure 1)**. In addition, SP-555 was co-localized with ACE2 (αACE2-488 (10 μg/ml) on the three cell types, but the hCMEC/D3 cells had a 2-fold greater colocalization compared to that of the other two cell types (**Figure 3A-F)**. Excess unlabeled αACE2 (60 μg/ml) reduced SP-555 uptake by about 40% for the hCMEC/D3 cells (**Figure 3G)**, which confirmed the data on SP-555 colocalization with αACE2-488 (10 μg/ml). Unlabeled excess SP (1 μM) displaced the bound labeled SP (SP-555) uptake by 50 to 65% for these cell types (**Figure 3H**). SP-555 uptake in the present of excess unlabeled SP is due to membrane bound and/or non-specific uptake. To establish the levels of extracellular receptor binding, SP-555 uptake was determined at 4 °C. At 37°C, SP-555 uptake was 2.2- to 5.5-fold greater than that at 4°C for these cell type. (**Figure 3I**). Thus, there was likely greater SP interaction with the cells and internalization (including membrane bound) at 37 °C. An illustration diagram of SP-555 uptake via ACE2 is shown in **Figure 3J**.

**Figure 3.**
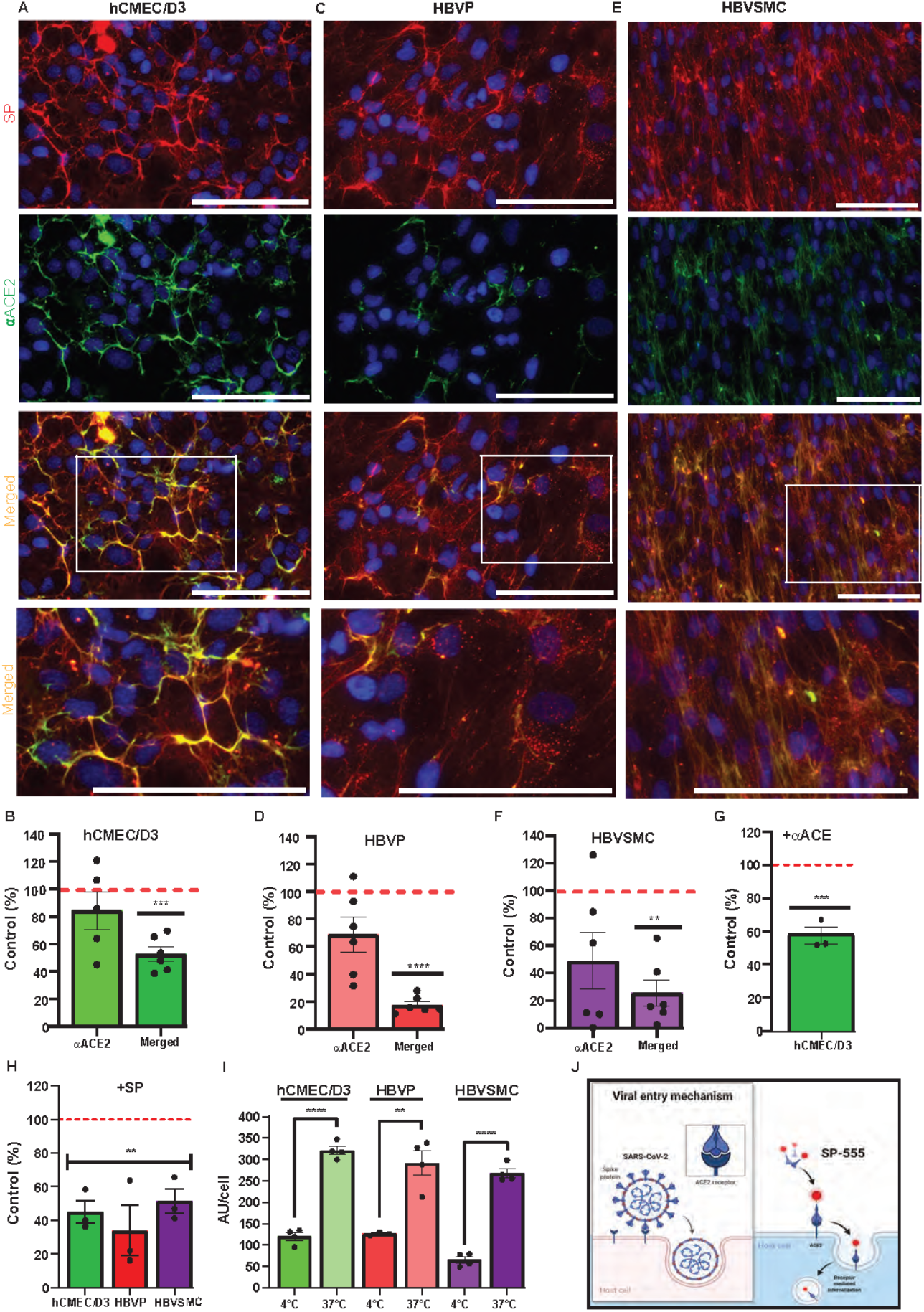
ACE2-mediated SP uptake. **A-F**. SP-555 and anti-human ACE2 antibody (αACE2-488) co-localized on the cell types, hCMEC/D3 (**A-B**), HBVP (**C-D**) and HBVSMC (**E-F**). Lowest row is the magnified image of the white boxed area above. Intensities of αACE2 and merged SP/αACE2 (yellow) were expressed as a percentage of SP-555 AU. **G**. Excess unlabeled αACE2 displaced SP-555 binding, which confirms the data in panel B. **H**. Excess unlabeled SP (self-competition) displaced SP-555 uptake. **I**. SP-555 receptor binding (4°C) considerable less than that at 37°C. **J**. Schematic diagram showing that SP-555 mimics SARS-CoV-2 binding to ACE2. Adapted from “Proposed Therapeutic Treatments for COVID-19 Targeting Viral Entry Mechanism” by BioRender.com (2021). Retrieved from https://app.biorender.com/biorender-templates. Red dashed line is the control levels (100%). Values are mean ± SEM. N= each data point is a well. Scale bar =100 μm. Supplementary Figure 1 and 3.

### Sialic acid/GM1-mediates SARS-CoV-2/SP uptake

Glycans, are carbohydrates based polysaccharides that are attached to molecules of the extracellular matrix of the cell surface, and can bind many toxins and pathogens, including viruses (Lingwood, 2011). Sialic acid containing glycan (N-acetyl D-glucosamine) has been reported to play a role in SP binding (Rhea et al., 2021). Similarly, glycosaminoglycans (GAGs), which are sulfate polysaccharides, are thought to play a role in infections (Aquino and Park, 2016; Dick and Vogt, 2014; Lima et al., 2017; Shi et al., 2021). While wheat germ agglutinin (WGA), a lectin that binds sialic acid, increased SP-555 uptake by all three cell types, it was 2.4- to 3.2-greater for hCMEC/D3 and HBVP cell types and 1.4-fold HBVMC (**Figure 4A-C**). SP uptake and internalization were increased with WGA (**Supplementary Figure 4**). In contrast, heparin, a polysaccharide, which belongs to the GAG family, did not affect SP uptake in the hCMEC/D3 cells but reduced its uptake to by 30 to 60% for HBVP and HBVSMC(**Figure 4A-C**). Anti-ganglioside (monosialotetrahexasylganglioside, GM1) antibody (αGMI) reduced SP-555 uptake by 60 to 80% of controls in all of these cell types (**Figure 4A-C**). However, in the presence of both antibodies (αACE2 and αGM1) the uptake was similar to that with αGM1 alone (**Figure 4D)**. Since GM1 is present in the lipid raft, SP binding by these cell types were caveolin/lipid raft-mediated. There are reports that SP can be taken up into cells by clathrin-mediated endocytosis and by binding to sialic acid residues on cell membrane bound glycoproteins (Bayati et al., 2021; Lingwood, 2011; Rhea et al., 2021). We confirmed that these cell types, including the endothelial cells, can take up transferrin (**Supplementary Figure 5A**), a molecule that is known to be transport by clathrin-mediated vesicles (Inoue et al., 2007; Mayle et al., 2012). BSA conjugated to lactosylceramide BODIPY, a molecule that is taken up by the lipid raft, was taken up by the cell membrane and colocalized with SP-555 (**Supplementary Figure 5B**). GM1-BODIPY was incorporated within the cell membrane (**Supplementary Figure 5C**). Thus, SP-555 is mainly cell membrane bound before internalization. While there is no specific inhibitor for each of these uptake mechanisms, nystatin (an inhibitor of lipid raft-mediated uptake) and chlorpromazine (an inhibitor of clathrin-mediated uptake) were tested (Plummer and Manchester, 2013; Sui et al., 2017; Wang et al., 2016). Both inhibitors were effective in blocking SP-555 uptake in each of these cell types by about 60%, except for chlorpromazine in HBVSMC, which inhibited it by 30% (**Figure 4E**). It’s tempting to speculate that SP uptake by GM1 (lipid raft) assist in its binding to ACE2 (**Figure 4F**), since both anti ACE2 and anti GM1 antibodies elicited the same effect as anti GM1 alone, and ACE2 is located within the caveolin/lipid raft (Garofalo et al., 2021; Glende et al., 2008; Lu et al., 2008).

**Figure 4.**
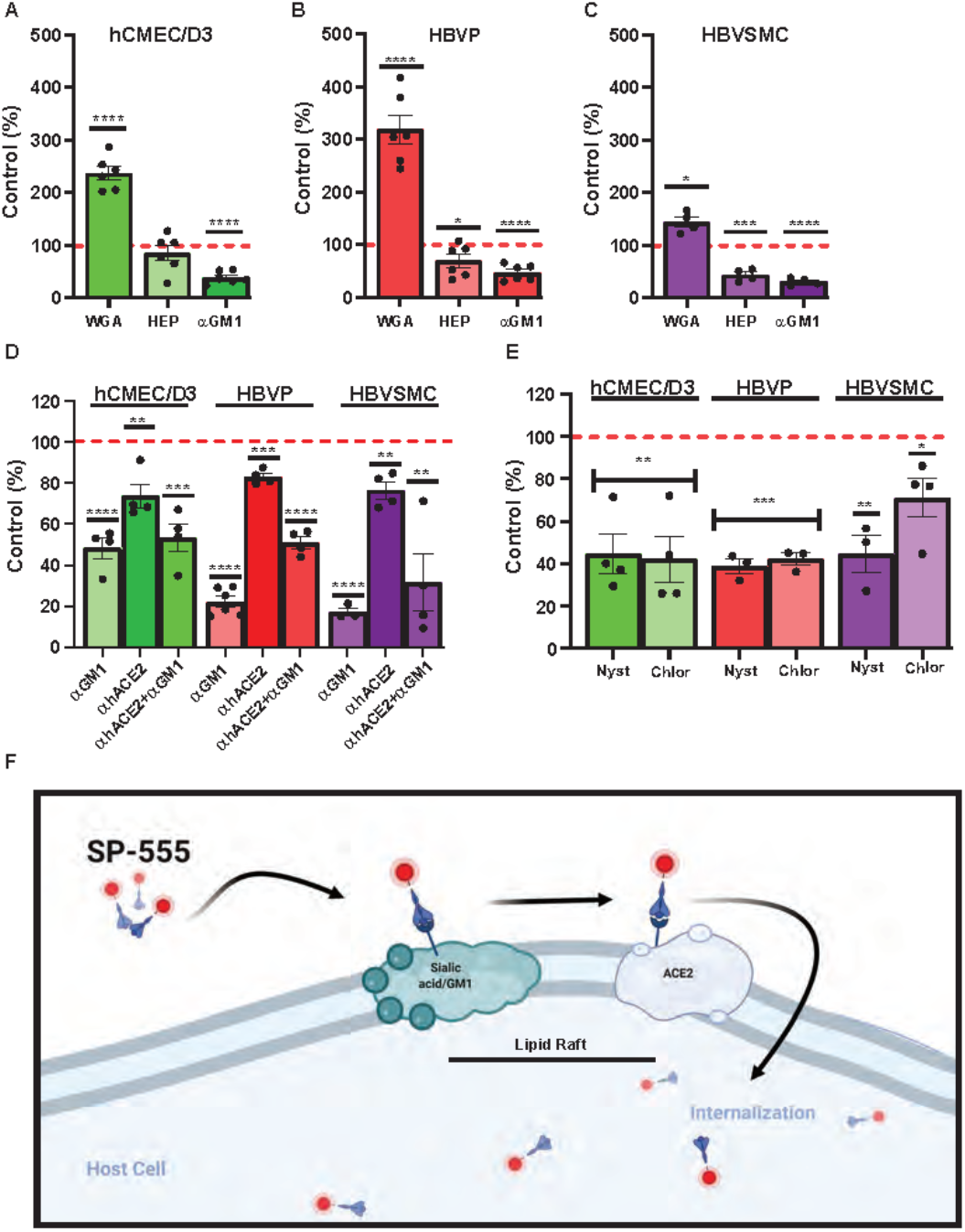
Sialic acid-/GM1-mediated SP uptake by cerebrovascular cells. **A-C**. Effect of wheat germ agglutinin (WGA), heparin (HEP) and anti-mono-sialotetra-hexasylganglioside antibody (αGMI) on SP-555 uptake. **D**. Effect of αGMI, anti-human ACE2 antibody (αACE2) and both αGMI and αACE2 together on SP-555 uptake. **E**. Effect of nystatin and chlorpromazine on SP-555 uptake. **F**. Schematic diagram showing a proposed SARS-CoV-2/SP uptake by both GM1 and ACE2. Both GM1 and ACE2 are present within the lipid raft region of cell membranes. Created by using BioRender.com. Red dashed line is the control levels (100%). Values are mean ± SEM. N=each data point is a well. Supplementary Figure 4

### Higher mutant SARS-CoV-2/SP uptake

While SARS-CoV-2 variants harbor many mutation sites, three mutation sites were selected from variants of concern (“WHO Coronavirus (COVID-19) Dashboard | WHO Coronavirus (COVID-19) Dashboard With Vaccination Data,” n.d.) to determine if their uptake is altered in these cell type (**Table 1**). These mutations were within the RBD (N501Y and E484K) and one (D614G), a common mutation site, which is outside the RBD and furin cleavage site (**Figure 5A**). For the hCMEC/D3 cells, the uptake of mutants D614G, N501Y and E484K was significantly increased by 1.5-, 1.9- and 2.8-fold, respectively, compared to control wild type SP (**Figure 5B**). In contrast, for the HBVP, the uptake of mutant D614G was unchanged, but for N501Y and E484K it was significantly increased by 1.7-fold compared to controls (**Figure 5C**). However, for the HBVSMC, the uptake of mutants D614G, N501Y and E484K was significantly increased by 3.2-, 5.0- and 3.8-fold, respectively, compared to controls (**Figure 5D**). Thus, again, there was differential mutant SP uptake by the three cell types. Uptakes of D614G, N501Y and E484K were highest for the HBVSMC, but there was no significant differences for the hCMEC/D3 and HBVP (**Supplementary Figure 6**). This may reflect the type and distribution of SP receptors. Mutation site E484K and N501Y confer gain-of-function, and N501Y increases SP affinity (Bayarri-Olmos et al., 2021; Tian et al., 2021; Xie et al., 2021). Mutant E484K also confers immune escape (Harvey et al., 2021; Weisblum et al., 2020). Mutant D614G seen to confers increased infectivity and transmissibility (Volz et al., 2021).

**Figure 5.**
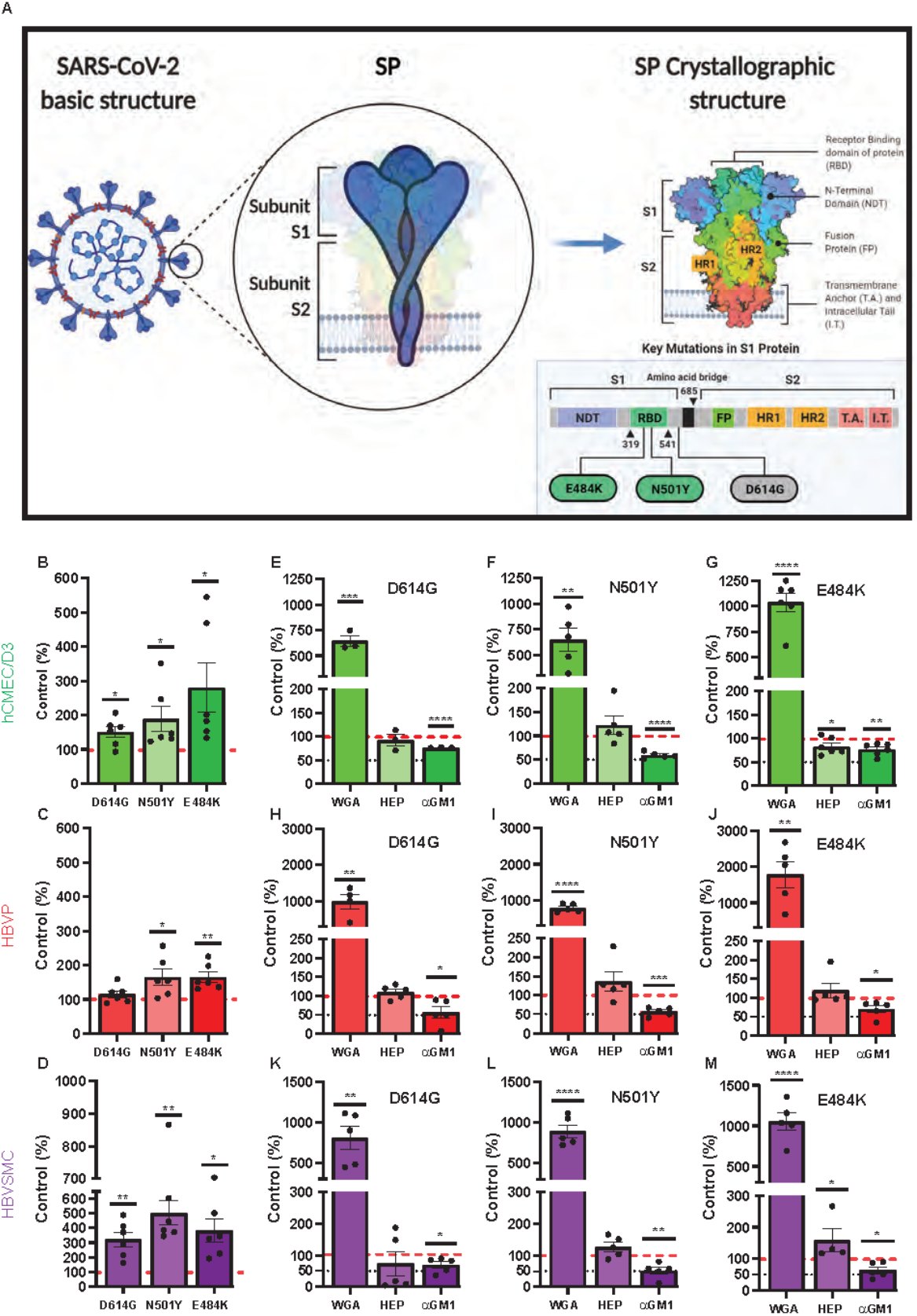
Increased mutant SP uptake. **A**. Schematic diagram showing the mutation sites of the three mutants (D614G, N501Y and E484K) used in this study. Adapted from “An In-depth look into the Structure of the SARS-CoV2 Spike Glycoprotein”, by BioRender.com (2021). Retrieved from https://app.biorender.com/biorender-templates. **B-D**. Mutant SP-555 uptake compared to control wild-type SP (red dashed line) for hCMEC/D3 (**B**), HBVP (**C**) and HBVSMC (**D**). **E-M**. The effects of wheat germ agglutinin (WGA), heparin and anti GM1 (a mono sialic acid ganglioside) on the three cell types, hCMEC/D3 (**E-G**), HBVP (**H-J**) and HBVSMC (**K-M**).Values are mean ± SEM. N= data points (wells) shown with the histogram. Red dashed line is the control levels (100%). Supplementary Figure 5.

Mutant SP uptake was considerably increased with WGA by 6.9- to 17.9-fold in the three cell types compared to their respective mutant control without WGA (**Figure 5E-M**). This was 2.9 to 7.7-fold greater than that seen for WGA with the wild type SP (**Figure 4A-C**). Mutant SP uptake was inhibited by anti-GM1 in the three cell types (**Figure 5 E-M**). In contrast, heparin had no effect except for E484K uptake in hCMEC/D3 and HBVSMC, which was decreased and increased, respectively (**Figure 5 E-M**). The neutralization effect of anti-GM1 antibody was less effective (**Figure 5 B-M**) by 1.5- to 3.1-fold compared to wild type SP (**Figure 4 A-C**). These differences may be due to the effectiveness of binding of the mutants SP and the distribution of the glycans and ACE2 on the three cell types. Excess αACE2 (60 μg/ml) suppressed SP E484K uptake (**Figure 6A-B**) but this was also less effective compared to wild type SP with the exception of hCMEC/D3 cells (**Figure 3A-G**).

**Figure 6.**
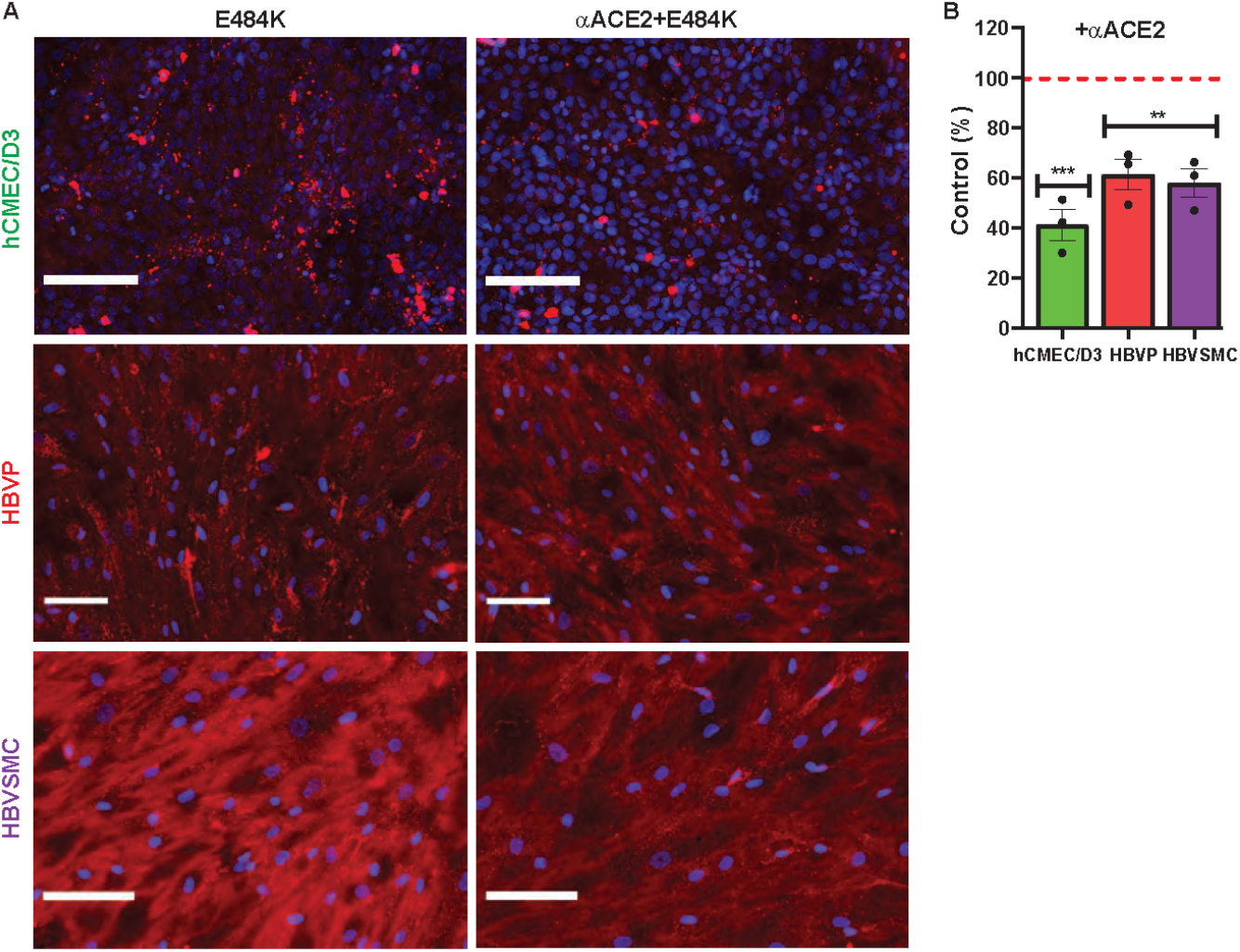
Anti-ACE2 suppressed mutant SP E484K uptake. **A-B**. Excess unlabeled αACE2 suppressed labeled SP E484K uptake in the hCMEC/D3 cells, HBVP and HBVSMC. Red dashed line is control levels (100%). Values are mean ± SEM. N= number of data points shown with the histogram. Scale bar =100 μm. Supplementary Figure 6

## Discussion

While there are neurological manifestations of COVID-19, it’s unclear if SARS-CoV-2 enters the normal human brain to contribute to this process (McQuaid et al., 2021). There are multiorgan tropism of SARS-CoV-2 which may indicate that the virus can be distributed to organs by blood(Brady et al., 2021). Establishing the mechanism of SARS-CoV-2 interaction at the cerebrovasculature will provide an understanding of its contribution to the neurological effects, which would facilitate the development of targeted therapies for specific symptoms.

While the endothelium is the physical site of the blood brain barrier (BBB), SARS-CoV-2 have to interact with other cell types, such as pericyte and smooth muscle cells to enter brain parenchyma. Since the SP on SARS-CoV-2 interacts with facilitators on the plasma membrane of host cell we used the SP to model viral behavior. We show that there was differential uptake of SARS-CoV-2/SP by these three human cerebrovascular cell types, endothelial cells, pericytes and vascular smooth muscle cells. The endothelial cell had the lowest capacity to take up SP compared to the other cell types, which may explain the low viral replication at the cerebrovasculature (Constant et al., 2021). In addition, there was saturable SARS-CoV-2/SP uptake, with similar Kd value for the endothelial and smooth muscles cells, while it was linear for pericytes. SARS-CoV-2/SP interact with ACE2, but this was mediated by GM1 as both of these are located within the caveolae/lipid raft. Glycosaminoglycans (GAGs) don’t seems to play a major role in SARS-CoV-2/SP uptake in endothelial cells, but plays a role in the pericyte and smooth muscle cells. WGA increased the SARS-CoV-2/SP uptake by each cell type.

Codons at position 484, 501 or 614 are common mutation sites in variants of concern, which are associated with increased binding to ACE2, transmissibility or immune escape (“WHO Coronavirus (COVID-19) Dashboard | WHO Coronavirus (COVID-19) Dashboard With Vaccination Data,” n.d.). There was increased N501Y, E484K and D614G uptake by all three cell types compared to wild type SP, except for D613G in pericyte. In the case of the endothelial cells, E484K uptake was increased by 2.8-fold compared to control SP. Uptake of the three mutants was increased 3.2-5.0-folds for smooth muscle cells. Thus, there is differential SP mutants by these cell types. A striking difference is that WGA considerable increased the binding of the three mutants SP by 6.9- to 17.8-fold, which was 2.9- to 7.3-fold greater than that of the wild type SP. Moreover, αACE2 and αGM1 were less effective in neutralizing mutant SP uptake compared to their effects on the wild type SP. In all cell types GM1, a glycosphingolipid ganglioside, contributes to SARS-CoV-2 binding to cerebrovascular cells. This glycolipid has diverse distribution and functions, which may contribute to many of the observed neurological symptoms in COVID-19.

Glycans, including gangliosides and glycosaminoglycans(GAGS), are carbohydrates based polymers that are attached to the extracellular surface of cell membrane, and can bind many toxins and pathogens, including viruses (Lingwood, 2011). This is a shield to protect cells (Nicoli et al., 2021; Sipione et al., 2020). GM1 is a glycosphingolipids ganglioside with one sialic acid containing oligosaccharide (glycan hydrophilic head) covalently bound to a ceramide lipid (hydrophilic tail). It is present in the ‘lipid rafts’ of the plasma membrane (Brown and London, 2000; D’Angelo et al., 2013; Degroote et al., 2004; Lingwood, 2011). It has diverse physiological functions, including neuroprotection, modulating cerebral blood flow and inflammation, mediate adhesion and protein recognition, cell signaling and communication, and membrane fluidity (Battistin et al., 1985; Furian et al., 2008; Hakomori, 2000; Iwabuchi et al., 1998; Jeyakumar et al., 2003; Mitsuda et al., 2002; Ngamukote et al., 2007; Vorbrodt, 1986). However, it also binds toxins, such as cholera and Shigo, and viruses, such as influenza and HIV (Chinnapen et al., 2007; Johannes, 2017; Suzuki, 2005) (Ilver et al., 2003; Lehmann et al., 2006; Varki, 2007). They are present at high levels on vascular endothelial cells, and involved in endocytosis and signaling (Born and Palinski, 1985; Weigel and Yik, 2002). Brain microvessels and cultured brain endothelial cells, including cell lines, express sialic acid oligosaccharides (N acetylglucosamine), which binds WGA (Fatehi et al., 1987; Plattner et al., 2010). SP binds to WGA and increased its uptake into brain, in mice (Erickson et al., 2021; Rhea et al., 2021). Our data confirm that WGA binds SP, but it’s mainly located on the endothelial cell surface, with little internalization. However, our data show that anti-GM1 antibody suppressed SP uptake in all three cerebrovascular cell type, and thus, a major facilitator for SP. Interaction of SP with GM1 could explain a recent study showing that SP caused an inflammatory response and reduced levels of tight junction proteins between endothelial cells using *in vitro* models, but a SP receptor was not identified in the report (Buzhdygan et al., 2020). Based on our data we proposed that GM1 interaction with SARS-CoV-2/SP contributes to the neurological manifestation of COVID-19.

Interestingly, Guillain-Barre syndrome (GBS) is associated with the presence of anti GM1 antibody, and there were reports of GBS-like effect in some SARS-CoV-2 vaccinated subjects (Kaida et al., 2009). GBS might be associated with vaccines against influenza and COVID-19 (Lunn et al., 2021; Martín Arias et al., 2015; Shapiro Ben David et al., 2021; Vellozzi et al., 2014). This raises intriguing questions on the role of cerebrovascular GM1 in COVID-19, including whether this has long term effects. It also offers a potential target to explore novel therapeutics to mitigate the ongoing perils of SARS-CoV-2.

Thus, SARS-CoV-2 could enter brain at the cerebravasculature, but this will be limited by the endothelial cells in normal healthy brains. It is likely that in the severely infected patients there is greater uptake by the brain due to the degree of cerebrovasculature dysfunction due to aging, prior health conditions, complication with the infections, and titer of the viral exposure. Cardio-respiratory failure is likely a major contributed to viral entry into brain.

### Limitations of the study

Since RBD of the SPs were used as a model of SARS-CoV-2, uptake of the actual virus might be different. However, the SP of SARS-CoV-2 is essential for viral entry into host cells. Thus, the attachment part of the viral life cycle can be explored with the SP. Also other cell types of the neurovascular unit, such as astrocytes, microglia and neurons need to be explored for SP uptake mechanisms. However, it is likely that there will be similar mechanisms since these cell types also express ACE2 and have glycans, including GM1.

## Contributors

CMQ performed experiments, imaging, preparation of figures and table and contributed to manuscript preparation; AS performed experiments, imaging, data analysis, preparation of figures and table, contributed to manuscript preparation and prepared the references; IA contributed resources and critical review of the MS; RD provide the concept, designed the study and wrote the manuscript. All authors approved the manuscripts.

## Acknowledgement

This work was supported by a NIH grant to RD (AG057574).

The Center for Musculoskeletal Research Histocore (CMSR) core facility (NIH P30 NIAMS AR069655). Schematic diagrams were created by using BioRender.com.

## Data and material availability

After publication the datasets used during the current study will be available from the corresponding author on reasonable request, and will be available in the Figshare repository

## Supplementary Information

### Title and legends

**Supplementary Figure 1. Human ACE2 binds SP555**. Recombinant human ACE2 dissolved in carbonate/bicarbonate buffer was immobilized (2 μg/ml) on glass slides for 1 hr at room temperature (RT), blocked with a non-protein buffer (Pierce Blocking buffer), washed, incubated with SP-555 at different concentrations in HBSS for 1 hr at RT, washed, mounted and imaged. SP-555 intensity from 10 fields for each concentration were analyzed and expressed as intensity/ μm^2^. Values are mean ± SEM.

**Supplementary Figure 2. SP not toxic to these cerebrovascular cells. A-C**. Cell viability determined by using the MTT cytotoxicity assay for the hCMEC/D3 (**A**), HBVP (**B**) and HBVSMC (**C**). Red dashed line is control levels (100%). Values are mean ± SEM. N=3 wells per group.

**Supplementary Figure 3. SARS-CoV-2 associated receptors present on these cerebrovascular cells. A-E**. Representative images (green) confirming the presence of ACE2, TMPRSS2, CD147, NP-1 and GM-1 on these cerebrovascular cell types (hCMEC/D3, HBVP and HBVSMC). Blue (DAPI) is the cell nucleus. Scale bar =50 μm.

**Supplementary Figure 4. Wheat germ agglutinin increased SP uptake**. Representative image of SP-555 uptake with and without wheat germ agglutinin (WGA) for hCMEC/D3. Scale bar =100 μm.

**Supplementary Figure 5. SP co-localized with ganglioside in these cerebrovascular cells. A**. Representative images confirming uptake of transferrin (TF-488). **B**. Representative images showing bovine serum albumin (BSA) conjugated to lactosylceramide BODIPY (BSA-BODIPY) uptake and its colocalization with SP-555. **C**. GM1-BODIPY uptake by these cells. Scale bar =100 μm.

**Supplementary Figure 6**.I**ncreased mutant SP uptake by the cerebrovascular cells. A-C**. Comparison of the mutants SP uptake between the three cell types (hCMEC/D3, HBVP and HBVSMC for D614G (**A**), N501Y(**B**) and E484K (**C**). Values are mean ± SEM. N= number of data points (wells) shown with each histogram. Red dashed line is the control levels (100%).

**Table 1. Main characteristics of SARS-CoV-2 variants containing the SP mutation sites used in this study**

