## Supplementary Figures for "SARS-CoV-2 spike proteins uptake mediated by lipid raft ganglioside GM1 in human cerebrovascular cells"

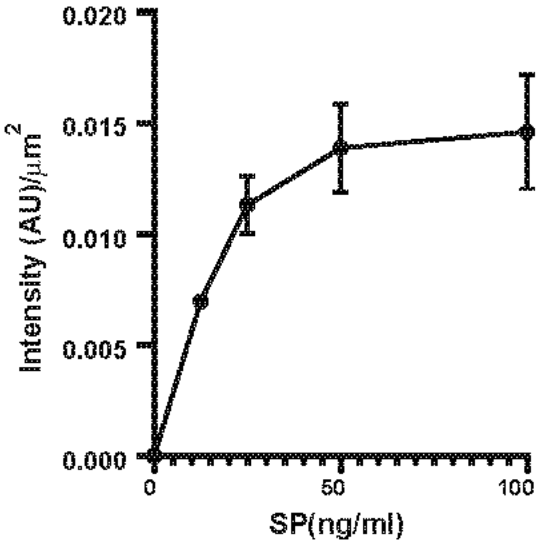


**Supplementary Figure 1. Human ACE2 binds SP555.** Recombinant human ACE2 dissolved in carbonate/bicarbonate buffer was immobilized (2 μg/ml) on glass slides for 1 hr at room temperature (RT), blocked with a non-protein buffer (Pierce Blocking buffer), washed, incubated with SP-555 at different concentrations in HBSS for 1 hr at RT, washed, mounted and imaged. SP-555 intensity from 10 fields for each concentration were analyzed and expressed as intensity/$\mu m^2$. Values are mean ± SEM.

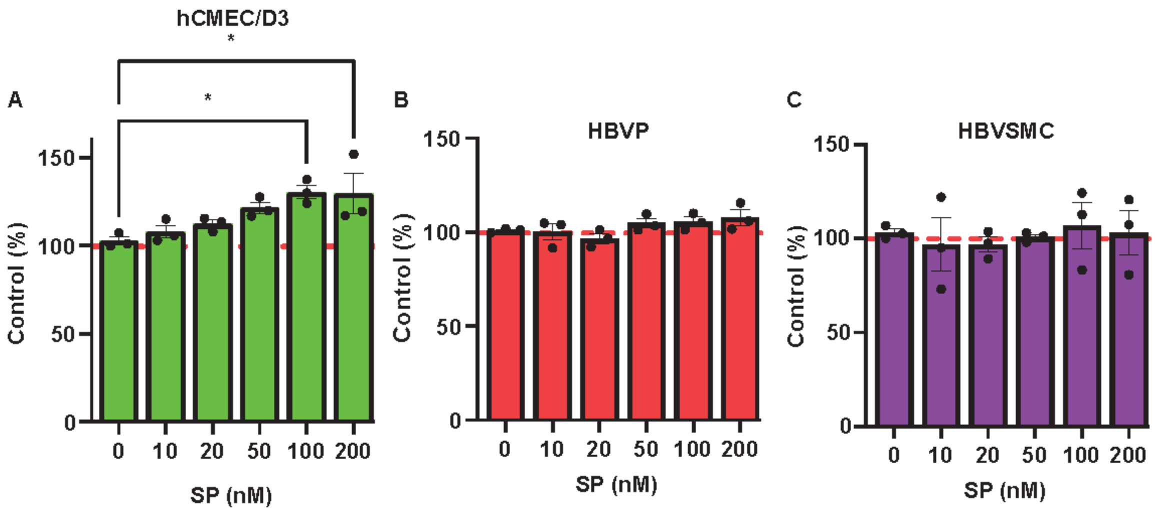


**Supplementary Figure 2**. **SP not toxic to these cerebrovascular cells. A-C**. Cell viability determined by using the MTT cytotoxicity assay for the hCMEC/D3 (**A**), HBVP (**B**) and HBVSMC (**C**). Red dashed line is control levels (100%). Values are mean ± SEM. N=3 wells per group.

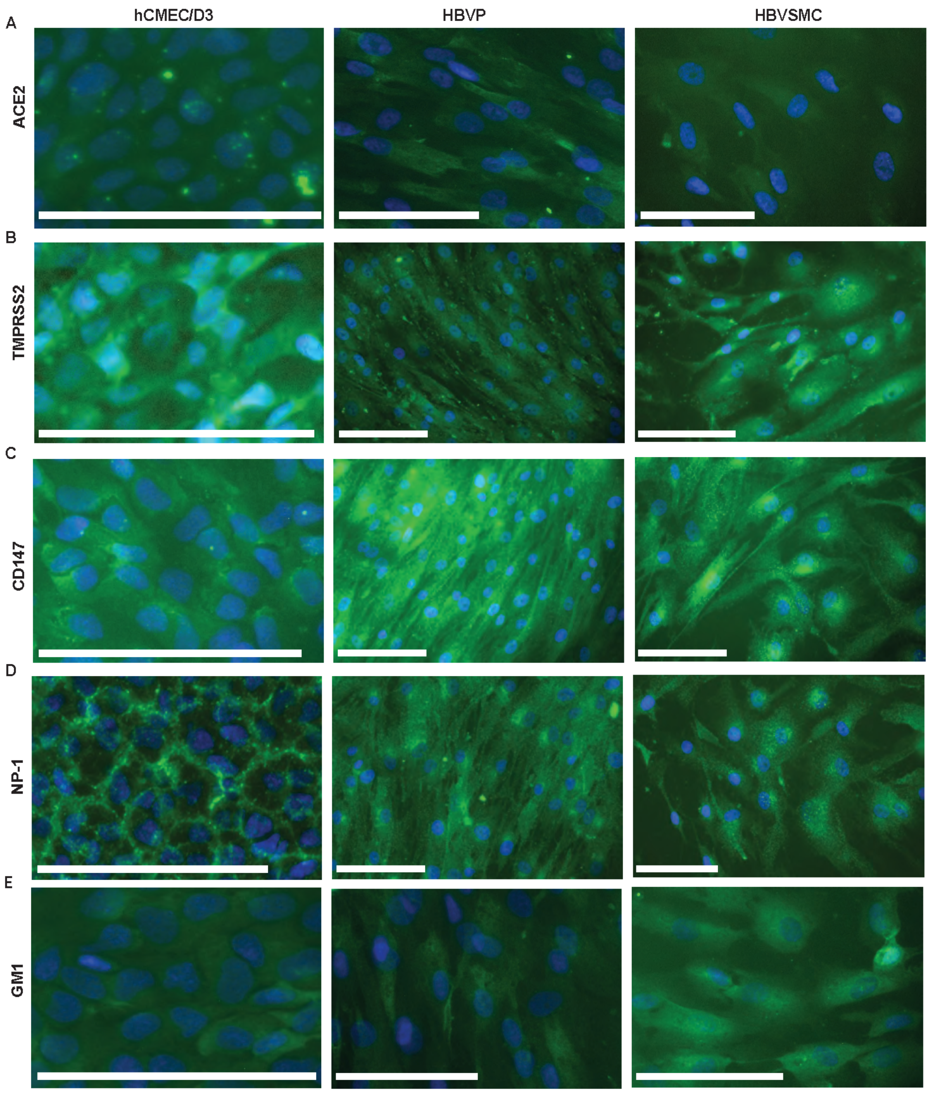


**Supplementary Figure 3**. **SARS-CoV-2 associated receptors present on these cerebrovascular cells**. **A-E**. Representative images (green) confirming the presence of ACE2, TMPRSS2, CD147, NP-1 and GM-1 on these cerebrovascular cell types (hCMEC/D3, HBVP and HBVSMC). Blue (DAPI) is the cell nucleus. Scale bar =50 μm.

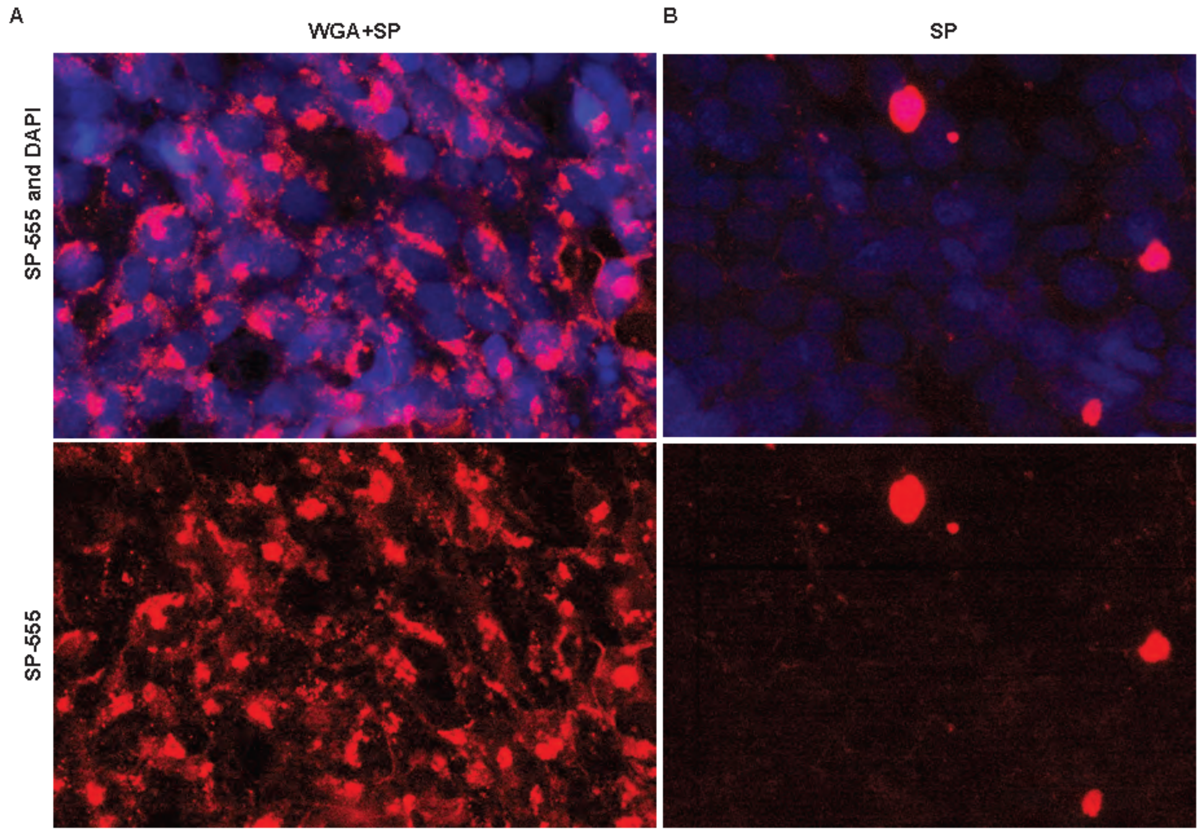


**Supplementary Figure 4. Wheat germ agglutinin increased SP uptake.** Representative image of SP-555 uptake with and without wheat germ agglutinin (WGA) for hCMEC/D3. Scale bar =100 μm.

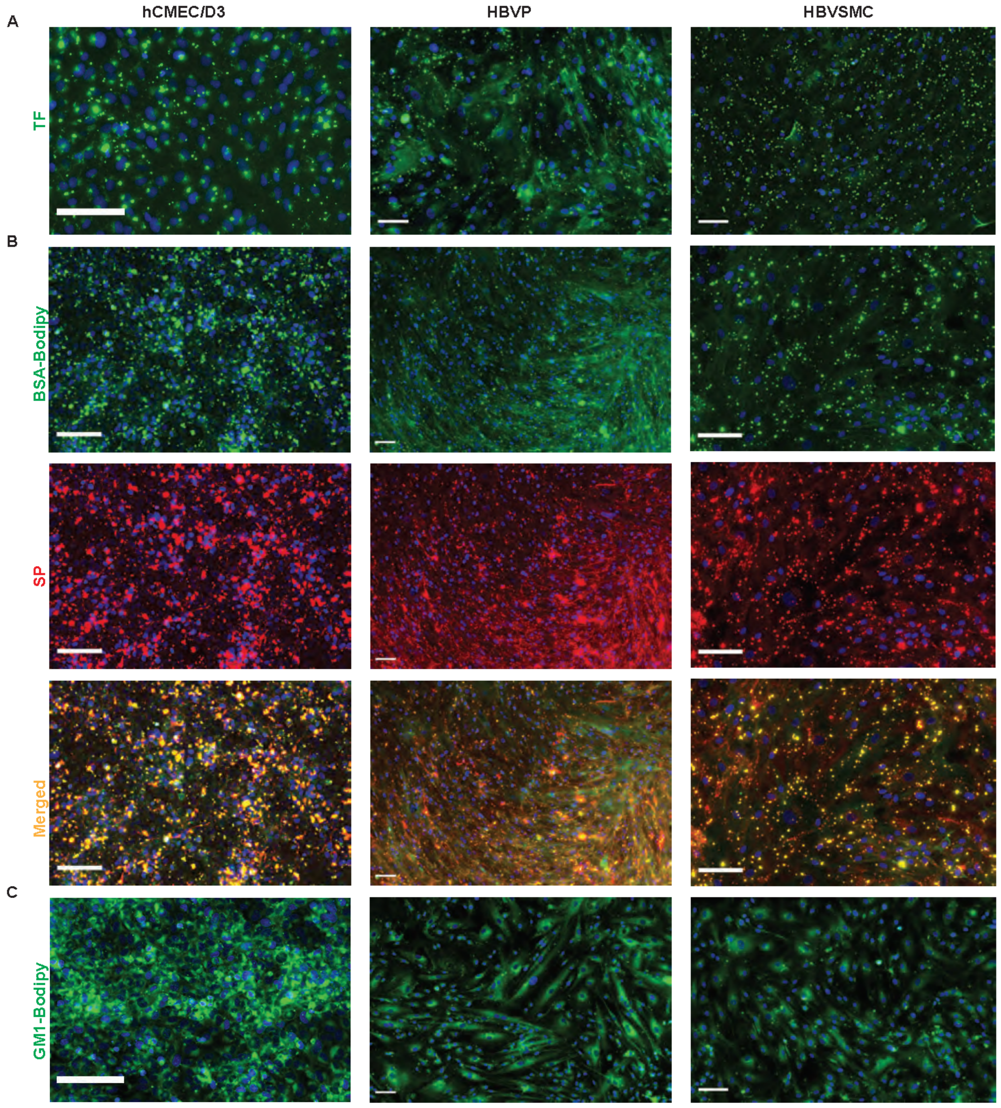


**Supplementary Figure 5**. **SP co-localized with ganglioside in these cerebrovascular cells**. **A**. Representative images confirming uptake of transferrin (TF-488). **B**. Representative images showing bovine serum albumin (BSA) conjugated to lactosylceramide BODIPY (BSA-BODIPY) uptake and its colocalization with SP-555. **C**. GM1-BODIPY uptake by these cells. Scale bar =100 μm.

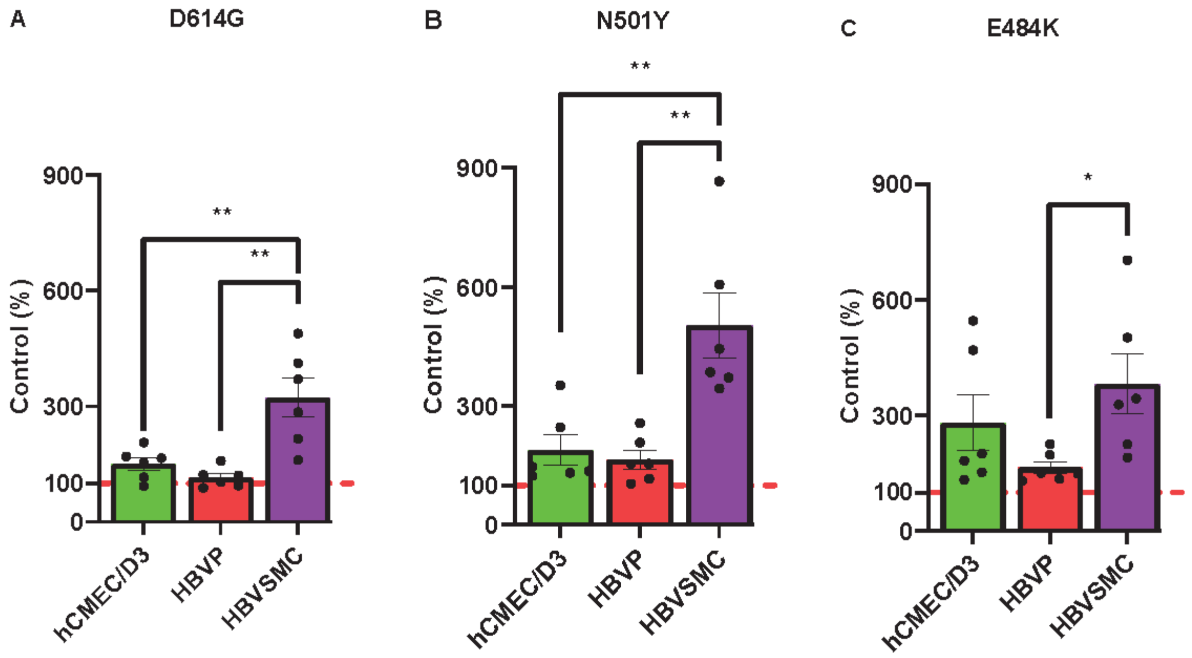


**Supplementary Figure 6.Increased mutant SP uptake by the cerebrovascular cells. A-C.** Comparison of the mutants SP uptake between the three cell types (hCMEC/D3, HBVP and HBVSMC for D614G (**A**), N501Y(**B**) and E484K (**C**). Values are mean ± SEM. N= number of data points (wells) shown with each histogram. Red dashed line is the control levels (100%).

### Table 1. Main characteristics of SARS-CoV-2 variants containing the SP mutation sites used in this study

| Variant of Concern | | Alpha | Beta | Gamma | Delta | Omicron |
|---|---|---|---|---|---|---|
| Pango Lineage(s) | | B.1.1.7 | B.1.351<br>B.1.351.2<br>B.1.351.3 | P.1<br>P.1.1<br>P.1.2 | B.1.617.2<br>AY. (1-12) | B.1.1.529.1<br>B.1.1.529.2<br>B.1.1.529.3 |
| Origin (date) | | UK (December 2020) | S.Africa (December 2020) | Brazil (November 2020) | India (October 2020) | S. Africa/Botswana (November 2021) |
| Critical mutation | RBD | N501Y | K417N, E484K, N501Y | K417T, E484K, N501Y | T478K, E484K, N501Y | E484A, T478K, N501Y |
| | Other | D614G, P681H | D614G | D614G | D614G, P681R | D614G, P681H, N679K |
| Main Properties | | -50% Increased transmission<br>-Potential increase severity<br>-Minimal impact on neutralization by convalescent and post vaccination sera | -50% increased transmission<br>-Reduced susceptibility to bamlanivimab and etesevimab<br>-Reduced neutralization by convalescent and post vaccination sera<br>-Increased severity | -Significantly reduced susceptibility to bamlanivimab and etesevimab treatments<br>-Reduction neutralization by convalescent and post-vaccination sera<br>-Significantly increased severity | -Increased transmissibility<br>-Potential reduction in neutralization by some antibody treatments<br>-Potential reduction by post-vaccination sera<br>-Increased severity | -Increased Transmissibility<br>-Reduced neutralization by some antibody treatments<br>-Less severe |

Omicron: Multiple countries of origin
